# The Madingley General Ecosystem Model predicts bushmeat yields, species extinction rates and ecosystem-level impacts of bushmeat harvesting

**DOI:** 10.1101/2020.03.02.959718

**Authors:** Tatsiana Barychka, Georgina M. Mace, Drew W. Purves

## Abstract

Traditional approaches to guiding decisions about harvesting bushmeat often employ single-species population dynamic models, which require species- and location-specific data, are missing ecological processes such as multi-trophic interactions, cannot represent multi-species harvesting, and cannot predict the broader ecosystem impacts of harvesting. In order to explore an alternative approach to devising sustainable harvesting strategies, we employ the Madingley General Ecosystem Model, which can simulate ecosystem dynamics in response to multi-species harvesting given nothing other than location-specific climate data. We used the model to examine yield, extinctions, and broader ecosystem impacts, for a range of harvesting intensities of duiker-sized ectothermic herbivores. Duiker antelope (such as *Cephalophus callipygus* and *Cephalophus dorsalis*) are the most heavily hunted species in sub-Saharan Africa, contributing 34%-95% of all bushmeat in the Congo Basin. Across a range of harvesting rates, the Madingley model gave estimates for optimal harvesting rate, and extinction rate, that were qualitatively and quantitatively similar to the estimates from single-species Beverton-Holt model. Predicted yields were somewhat greater (around 5 times, on average) for the Madingley model, which would be expected given that the Madingley simulates multi-species harvesting from an initially pristine ecosystem. This match increased the degree of confidence with which we could examine other predictions from the ecosystem model, as follows. At medium and high levels of harvesting of duiker-sized herbivores, there were statistically significant, but moderate, reductions in the densities of the targeted functional group; increases in small-bodied herbivores; decreases in large-bodied carnivores; and minimal ecosystem-level impacts overall. The results suggest that general ecosystem models such as the Madingley model could be used more widely to help estimate sustainable harvesting rates, bushmeat yields and broader ecosystem impacts across different locations and target species.

## 1. Introduction

Present levels of wild animal harvesting are believed to be a major threat to survival for over half of the 178 species currently hunted in Central Africa (Abernethy *et al.*, 2013). Bushmeat harvesting is an essential source of food and income for many poor rural communities in sub-Saharan Africa (Milner-Gulland and Bennett, 2003; Davies and Brown, 2008; Fa, Currie and Meeuwig, 2003). Declining animal abundances and potential loss of species will detrimentally affect biological diversity and ecosystem integrity (Abernethy *et al.*, 2013; Hooper *et al.*, 2005), as well as the livelihoods and wellbeing of human population relying on meat from wild animals (or bushmeat) for cash income and additional protein (Nasi, Taber and Van Vliet, 2011; Golden *et al.*, 2011; Njiforti, 1996; Foerster *et al.*, 2011).

The standard approach to modelling the impacts of harvesting on a wild population is to use a population dynamic model, parameterised for the target species. A limitation is that single-species models are limited to studying the impacts of harvesting on that single species, and require species-specific or location-based data, and parameter estimates. By contrast, a practical approach to bushmeat harvesting over whole regions will require methods that can estimate the impacts of harvesting multiple species (Fa and Peres, 2001), on both the target and the non-target species (Abernethy *et al.*, 2013), over large regions where species- and location-specific data are sparse or not available (Fa and Brown, 2009).

The modelling approaches currently used for assessing sustainability of bushmeat harvesting rely heavily on species monitoring data. These methods involve examining changes in animal abundances (e.g. Van Vliet *et al.*, 2007) and harvest offtakes over time (e.g. Albrechtsen *et al.*, 2007). Although declines in abundances of targeted species have been attributed to overharvesting in a number of Central African study sites, observational data is generally too limited (temporally, spatially) and/or too variable to reliably inform an effective management strategy (Wilkie *et al.*, 2001; Linder, 2008; Gates, 1996).

Instead of using time-series data on animal densities and offtakes, sustainability indices, such as Robinson and Redford’s index (Redford and Robinson, 1991) could be used to estimate sustainable harvest rates. This allows an estimation of sustainable levels of production of harvested populations (Wilkie and Carpenter, 1999; van Vliet and Nasi, 2008; Fa *et al.*, 2014) which can then be compared with actual data on animal offtakes. However, once again, to be effective most sustainability indices require accurate estimates of population parameters (Milner-Gulland and Akçakaya, 2001; van Vliet and Nasi, 2008; Weinbaum *et al.*, 2013), such as the population carrying capacity and rate of population growth.

In practice, multiple species are targeted by hunters in tropical forests. To-date, optimising harvesting beyond a single-species approach has been studied in theory (Bhattacharya and Begum, 1996; Song and Chen, 2001) and attempted in fisheries management (Yodzis, 1994; Hutniczak, 2015), where multi-trophic relationships are better described than in terrestrial ecosystems. Attempts to combine the understanding of multi-trophic interactions, current knowledge of biophysical systems (climate, nutrient flows, ecological processes) and how humans interact with the system (offtake levels, monitoring, socioeconomic drivers of demand) resulted in a number of ecosystem models for separate biomes (Goodall, 1975; Travers *et al.*, 2007; Metzgar *et al.*, 2013); but none of the terrestrial ecosystem models have been used for decision-making in practice. More recently, sophisticated end-to-end marine ecosystem models, such as Atlantis (Fulton *et al.*, 2004, 2011) and Ecopath with Ecosim (EwE) (Christensen and Walters, 2004) have been developed and have now been applied to many marine ecosystems (for example, about 130 EwE models have been published, Travers *et al.*, 2007). However, deployment of these models requires extensive data inputs such as place-specific biological parameters (e.g. production rate, diet composition) and stock assessment survey data for a number of selected functional groups (Link, Fulton and Gamble, 2010; Travers *et al.*, 2007). Consequently, these modelling frameworks cannot be applied without extensive parameterisation and good knowledge of the system (Link, Fulton and Gamble, 2010).

New datasets including ones on global animal density (TetraDENSITY; Santini, Isaac, and Ficetola 2018), biodiversity (PREDICTS; Hudson *et al.*, 2017) and bushmeat harvesting (Offtake; Taylor *et al.*, 2015) have been developed, and new computational methods (e.g. Bayesian and Machine Learning) could be used to make the most of these new data. However, despite these efforts, the extent (taxonomic, spatial, temporal) of species-level data in sub-Saharan Africa is still very limited, especially in the regions where bushmeat harvesting is of highest concern (e.g. sub-Saharan Africa) where there are no data at all available for the vast majority of the harvested species (Rodríguez *et al.*, 2007; Fa and Brown, 2009).

In terms of the effects of harvesting on ecosystem structure and functioning, a number of studies have reported increases in non-target species abundances (Peres and Dolman, 2000; Linder, 2008). Peres (2000) showed that species resilience to harvesting correlated with body size (large-bodied species were more sensitive to persistent harvesting) in the Amazonian tropical forests. However, bushmeat harvesting studies in tropical forests generally focus on impacts of harvesting on the target species.

Thus, new methods are needed that deal with data scarcity while reflecting uncertainty and that incorporate ecosystem and community impacts (Weinbaum *et al.*, 2013). One potential solution is to employ an ecosystem model whereby fundamental ecological principles are used to simulate ecosystem structure and function, allowing emergent macroecological patterns to develop bottom-up. The Madingley General Ecosystem Model (hereafter referred to as the Madingley model) is such a model and has been tested against observations in a variety of virtual experiments (Harfoot *et al*., 2014). It can simulate the effects of alternative harvesting scenarios on all species in the ecosystem, without the need for location- or species-specific data or parameters. It therefore offers an alternative to the traditional data-driven approaches currently in use in terrestrial harvesting.

To date, the Madingley model (Purves *et al.*, 2013; Harfoot *et al.*, 2014) is the only such mechanistic ecosystem model that can be applied to any ecosystem type (marine and terrestrial), at any location, and at any spatial resolution level. It shares some important features with other ecosystem models such as aggregation of organisms into functional groups and the inclusion of biophysical drivers (climate, net primary production). However, unlike analogous models, the aggregation is not species-specific: it takes place on a functional level, based on traits such as diet (herbivore, carnivore, omnivore), metabolism (warm vs cold blooded) and adult and current body size, all of which are treated with well-established ecological relationships. Ecosystem dynamics (animal and plant) emerge in the Madingley model as a result of environmental inputs (such as air temperature and precipitation levels) working upon animals and plants, whose interactions between themselves and with the environment are based on fundamental concepts and processes derived from ecological theory, and defined at the scale of the individual organism. Importantly, all of these details mean that the model can simulate ecosystem dynamics at any location, without the need for explicit parameterisation by species or location. All that needs to be specified is the location (latitude, longitude) because this is needed to look up the climate drivers; and any perturbations made to the system. Crucially for this paper, these perturbations could include harvesting of any combination of plants and animals from the system.

On a functional group level, the Madingley model has been shown to provide robust approximations of the dynamics of animal populations (Harfoot *et al.*, 2014). The model’s outputs are spatially explicit, and allow for the calculation of whole-ecosystem metrics such as animal abundance, body mass and trophic indices, which could all be used as indicators of systems’ sensitivity to perturbations. To date, the Madingley model is the only, to our knowledge, model allowing such ecosystem-wide questions to be explored without specific and detailed parameterisation.

Here, we run a series of experiments in the Madingley model to compare the estimates it provides of sustainable harvesting bushmeat harvesting from an African tropical rainforest. Duiker (*Cephalophinae*) is the most heavily hunted species in sub-Saharan Africa contributing 34%-95% of all bushmeat captured in the Congo Basin (Wilkie and Carpenter, 1999; Fa, Ryan and Bell, 2005). Unusually, duikers have at least some species-specific ecological data, which allows us to compare the Madingley model predictions to predictions from species-specific single-species models. We then examine the Madingley model predictions for broader ecosystem impacts of harvesting, which cannot be done with the single-species models. We are interested in the model’s estimates of sustainable harvesting in the tropical forest ecosystem, and the potential impacts of harvesting on ecosystem structure. We are ultimately interested in whether such an approach, using ecosystem modelling, could be developed to be useful in practice for the vast majority of the worlds harvested species and locations which, unlike African duikers, have not been surveyed at all.

## 2. Methods

We begin by running the Madingley simulations for harvesting duiker *Cephalophus* spp. We create a Madingley model experiment that is as close as possible to those already run in Barychka *et al*. (in prep.) using the single-species model (Beverton-Holt; Beverton and Holt, 1957), to allow comparison of the outputs. The single-species model is parameterised using empirical estimates for Peters’ duiker *C.callipygus* and bay duiker *C.dorsalis* (Feer, 1988; Lahm, 1993; Fa *et al.*, 1995; Noss, 1998a; Noss, 1998b; Hart, 2000; Noss, 2000; van Vliet and Nasi, 2008), so qualitative and/or large (higher than first order; Coe, Cumming and Phillipson, 1976) quantitative differences between the models’ outputs would increase our level of scepticism about using the Madingley model. On the other hand, good level of correspondence between the models would increase our level of confidence in examining the Madingley predictions that the single-species model cannot make. Hence, we view this as a ‘validation experiment’. We look closely at the yield, and the maximum harvest rate, for duikers as predicted by the Madingley model, including reporting on the uncertainty in the yields (see 3.1). This much was possible using the single-species model. However, we also examine the impact of duiker-like harvesting on the structure of the whole ecosystem (see 3.2), something that is only possible with the Madingley model. This allows us to assess whether and how apparently sustainable harvesting, could affect ecosystem structure.

### 2.1 Simulation Protocol

#### The models

A schematic representation of the Madingley model (with harvesting) is given in Figure 1, along with a representation of a single-species model (with harvesting). The Madingley model: a) receives environmental data based on user-defined latitude and longitude: location-specific empirical data on air temperature, precipitation levels, number of frost days, seasonality of primary productivity and soil water availability; b) simulates ecosystem dynamics from environmental inputs, and animal and plant dynamics described in the model using a set of core biological and ecological functional relationships (plant growth and mortality, and eating, metabolism, growth, reproduction, dispersal, and mortality for animals); and c) outputs estimates of biological characteristics of the emergent ecosystem (Harfoot *et al.*, 2014).

**Figure 1.**
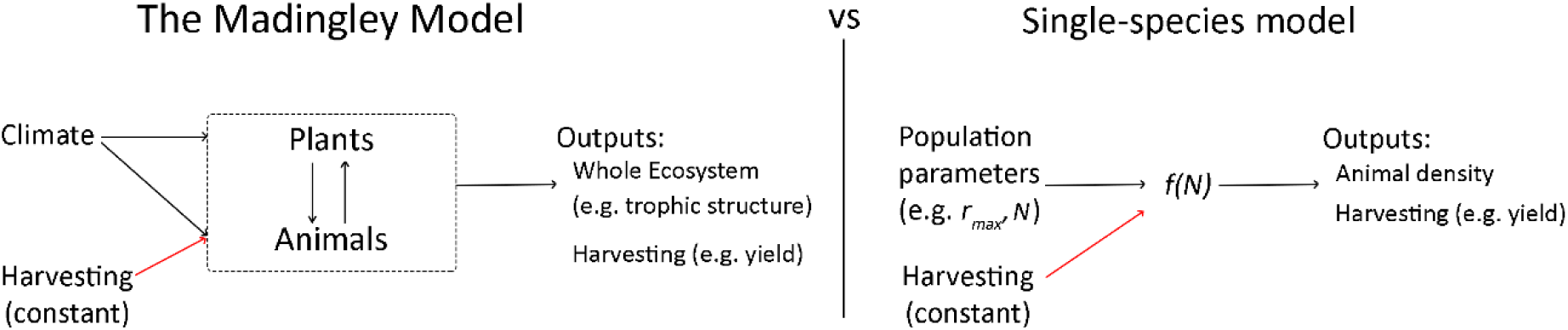
The Madingley model’s inputs, modelled processes and outputs, compared to a single-species model’s inputs, processes and outputs.

The Madingley model represents the state of the consumer (animal) part of the ecosystem in terms of the densities of individual animals with different functional traits. The densities change through time as individuals interact, in turn resulting in births, deaths, growth, and dispersal, with the interactions (e.g. predation) defined entirely in terms of those traits. Although the model is defined by interactions among individuals, the simulation uses a computational approximation (based around so-called cohorts) to allow for all interactions among all individuals to be simulated. The animal part of the ecosystem is ultimately fed by the vegetation, which growth is simulated using a simple stock and flow model, driven by climate, but affected by herbivory. For detailed description see Harfoot *et al.*, (2014).

As a comparison for the Madingley predictions, we used the Beverton-Holt population dynamics model (Beverton and Holt, 1957) to represent single species responses to harvesting pressure:

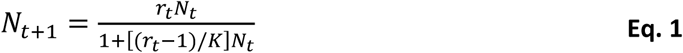

where *N*_*t*_ is the population density (individuals per unit area: in this case, animals km^-2^) at time *t*; *N*_*t*+1_ is the population density in the following time step; *K* is the equilibrium population size in the absence of harvesting; and *r* = *exp*(*r*_*max*_) is the density-independent intrinsic rate of natural increase (the balance of births and deaths) for year *t*.

The Beverton-Holt model has been widely used in the past to study the dynamics of harvested species (e.g. Barnes, 2002; Holden and Conrad, 2015); it is compensatory rather than over-compensatory (high density leads to a reduction in per capita reproduction but does not reduce the recruitment of the entire population; Kot, 2001) and is believed to provide a good representation of intraspecies competition in ungulate populations that are not constrained by resources or habitat availability (Ruckstuhl and Neuhaus, 2000).

#### Location

Our experimental site was a simulated on a 1^0^ × 1^0^ geographic grid cell (111.32km x 110.57km) centred on 1°S, 15^0^E; the coordinates were selected to fall within the known duiker range in the tropical forests of the Republic of Congo. For the purposes of this study, no inter-cell migration was modelled, i.e. no animals were allowed from outside the experimental area.

#### Target group

We simulated harvesting strategies for animals similar to duiker antelope (Table 1). We set up harvesting in the Madingley model to target terrestrial herbivorous endotherms, described using the following categorical traits: ‘Heterotroph - Herbivore - Terrestrial - Mobile - Iteroparous - Endotherm’. This definition was further narrowed using two continuous traits: adult body mass and juvenile body mass (Lahm, 1993; Noss, 1998a). Under this definition, the target group for duiker-like harvesting included two out of the three most heavily hunted duiker species in Central Africa (Noss, 1998a): Peters’ duiker *Cephalophus callipygus* and bay duiker *Cephalophus dorsalis*. This excluded smaller-bodied herbivores (such as blue duiker *Cephalophus monticola*), but also other bushmeat species such as medium-sized herbivorous primates (such as *Piliocolobus badius*, mean weight = 7.75kg, mean density = 156.3 animals/km^2^) and large rodents (such as *Thryonomys swinderianus*, mean weight=5.05kg; mean density=9.97 animals/km^2^) (Fa, Ryan and Bell 2005).

**Table 1.** Summary of harvesting experiment in the Madingley model: harvesting duiker-like herbivores (13-21kg). We reduced the size of the steps for harvest rates of 0.25-0.60 to examine the model’s outputs and dynamics in more detail around the optimum harvest rates.

| Target group | Madingley traits | Harvest rate, $\phi$ | Example species | Response metrics |
| --- | --- | --- | --- | --- |
| Duiker-like: 13-21kg as adults and >100g as juveniles | Endothermic Herbivores | 0.00-0.25 in steps of 0.05 and<br>0.25-0.60 in steps of 0.03 and<br>0.60-0.90 in steps of 0.10 | Peters' duiker *Cephalophus callipygus*;<br>Bay duiker *Cephalophus dorsalis* | • Yields (animals km<sup>-2</sup> year<sup>-1</sup>);<br>• Survival Probability (over 30 years);<br>• Change in Ecosystem Structure |

#### Harvesting

In the Madingley model, a 1000-year ‘burn-in’ (no-harvesting) period was run (*n*=30) to produce estimates of the ecosystem’s equilibrium state in year 1000, including, for each functional group (carnivore/omnivore/herbivore): the number of surviving animal cohorts, abundances, biomass, and adult body masses. These estimates of ecosystem’s equilibrium ecological community were used as a starting point for subsequent harvesting simulations (i.e. the same 30 burn-in simulations were used as inputs for the subsequent harvesting simulations).

We used a constant proportional harvesting policy (Case, 2000), where each year a proportion (harvest rate *φ*, Table 1) of animals were targeted. This harvest rate remained constant for the duration of harvesting period *t* (set at 30 years based on examining outputs’ sensitivity to harvesting duration, results not shown here). Experiments were replicated 30 times at each harvest rate: a larger sample size of 100 was also attempted for a selection of harvest rates; however, resulting dynamics did not differ significantly from a smaller sample of 30, and the time needed to run the simulations was substantially higher. Harvesting took place once a year in month 6: we simulated discrete harvesting (as opposed to continuous) to better approximate harvesting in the Beverton-Holt model.

In the Beverton-Holt model, simulations were run following methodology in Barychka *et al*. (in prep.). Parameters *r*_*max*_ and *K* were derived from field observations and included uncertainty. We simulated proportional harvesting over 30 years with harvest rate *φ* ranging from 0 (no harvest) to 0.90 in discrete steps of 0.05, giving 19 different values of *φ*. For each combination of timescale and harvest rate, we carried out an ensemble of 1000 simulations. Harvesting was applied from year 1 onwards (no harvesting took place in year 0). The ensemble size was based on preliminary analysis involving comparing summary statistics and visualising results for smaller (100 simulations and 500 simulations) and larger (10000 simulations) sample sizes. Based on model estimates, we assessed average yields, survival probability, and the uncertainty in both yield and survival.

### 2.2 Output Metrics

#### Yield

Using the Madingley model, total yields and target animal densities were recorded. The total yield in year *t* was equal to 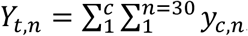, where *y*_*c,n*_ was yield from harvesting cohort *c* in simulation *n* in month 6. The total density was *D*_*m,n*_ = Σ*d*_*m,c,n*_, where *d*_*m,c,n*_ was density for target cohort *c* in simulation *n* in month *m*.

Using the Beverton-Holt model, yield at time *t* (*Y*_*t*_) was the difference between the number of animals at time *t* (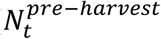 after reproduction at the end of year *t* − 1), and the higher of 0 and the number of surviving animals after target proportion *φ* of animals had been extracted at time *t*. I.e. 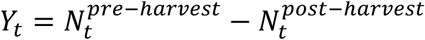, where the number of animals that remain in the population after harvesting at time *t*, 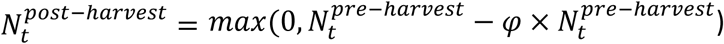.

#### Extinction

The rate of extinction of duiker-like animals in the Madingley model was estimated at each time step. Extinction was defined when the total density (*D*_*m,n*_) of animals that matched the definition of duiker-like fell below 0.1 animals km^-2^ during a simulation run. This corresponds to approximately 99% reduction in density from average carrying capacity for Peters’ and bay duiker (Feer, 1988; Lahm, 1993; van Vliet and Nasi, 2008).

The same threshold (0.1 animals km^-2^) was applied to estimate the two duiker species survival probability using the Beverton-Holt model. A response of 1 was assigned to a year where population size 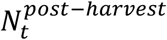 was equal to or above a threshold of 0.1 animals km^2^; zero (0) was assigned to a year (and all subsequent years) when population size dipped below the viability threshold (we set 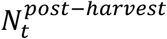, i.e. quasi-extinction). Responses were then averaged to give an estimate of survival probability at each harvest rate with 95% confidence intervals over 30-year harvest.

#### Ecosystem Response

The ecosystem-level information was recorded in the Madingley at each time step, such as, for each functional group, adult body masses, animal biomasses and abundances.

Overall, the ecosystem-level response to harvesting was analysed as follows. First, each cohort was identified to functional group (*f*) as being a herbivore, omnivore or carnivore. Individuals were also allocated into a body mass bin (*b*) ranging from the smallest body mass (10^−2^ to 10^−1^ gram; *b* = −2) to the largest bin (10^6^ to 10^7^ grams; *b* = 6). Because some of the bins were deemed too wide to be able to capture changes in cohort abundances due to harvesting (see Figure S1), bins were further sub-divided into smaller sub-bins, where adult body masses were incremented in steps of 0.5 for 2 ≤ *b* ≤ 6 (Figure 3 and Figure S2a), and in even smaller increments of 0.25 for 3 ≤ *b* ≤ 5 (Figure S2b). Total abundances were then calculated for each functional group in each body mass bin, logged (on log10 scale) and normalised to month 1 of the simulation for visualisation purposes.

**Figure 2.**
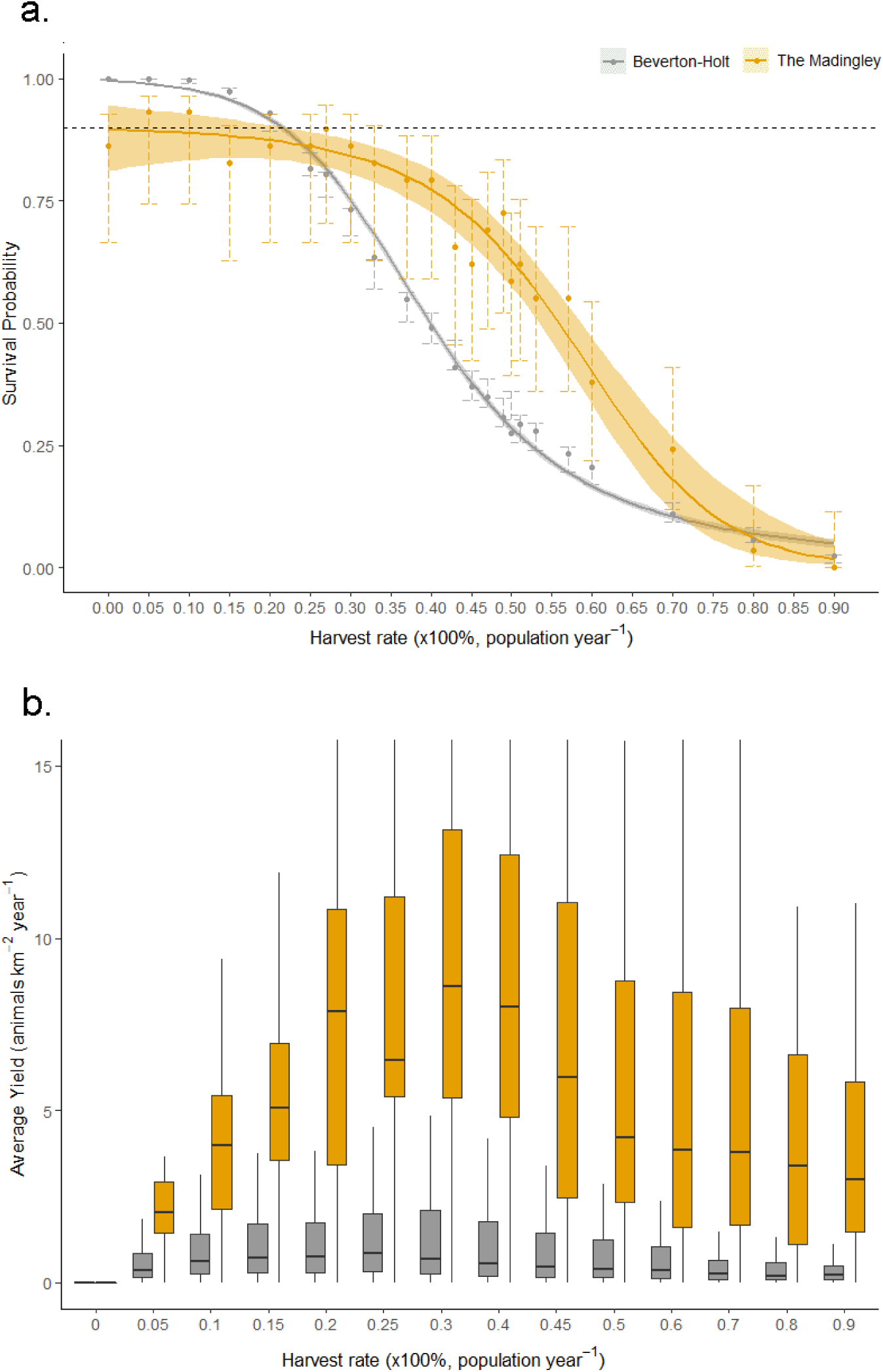
Survival probabilities (with 95% confidence intervals in grey/orange shading and 2 standard errors indicated by vertical error bars) in a. and estimated yields in b., from proportional harvesting of Peters’ duiker using the Beverton-Holt model (in grey), and of duiker-like herbivores (13-21kg) using the Madingley General Ecosystem Model (in orange). The horizontal dashed line in a. indicates a 90% survival target (i.e. extinction in less than 10% of the cases; Mace and Lande, 1991).

**Figure 3.**
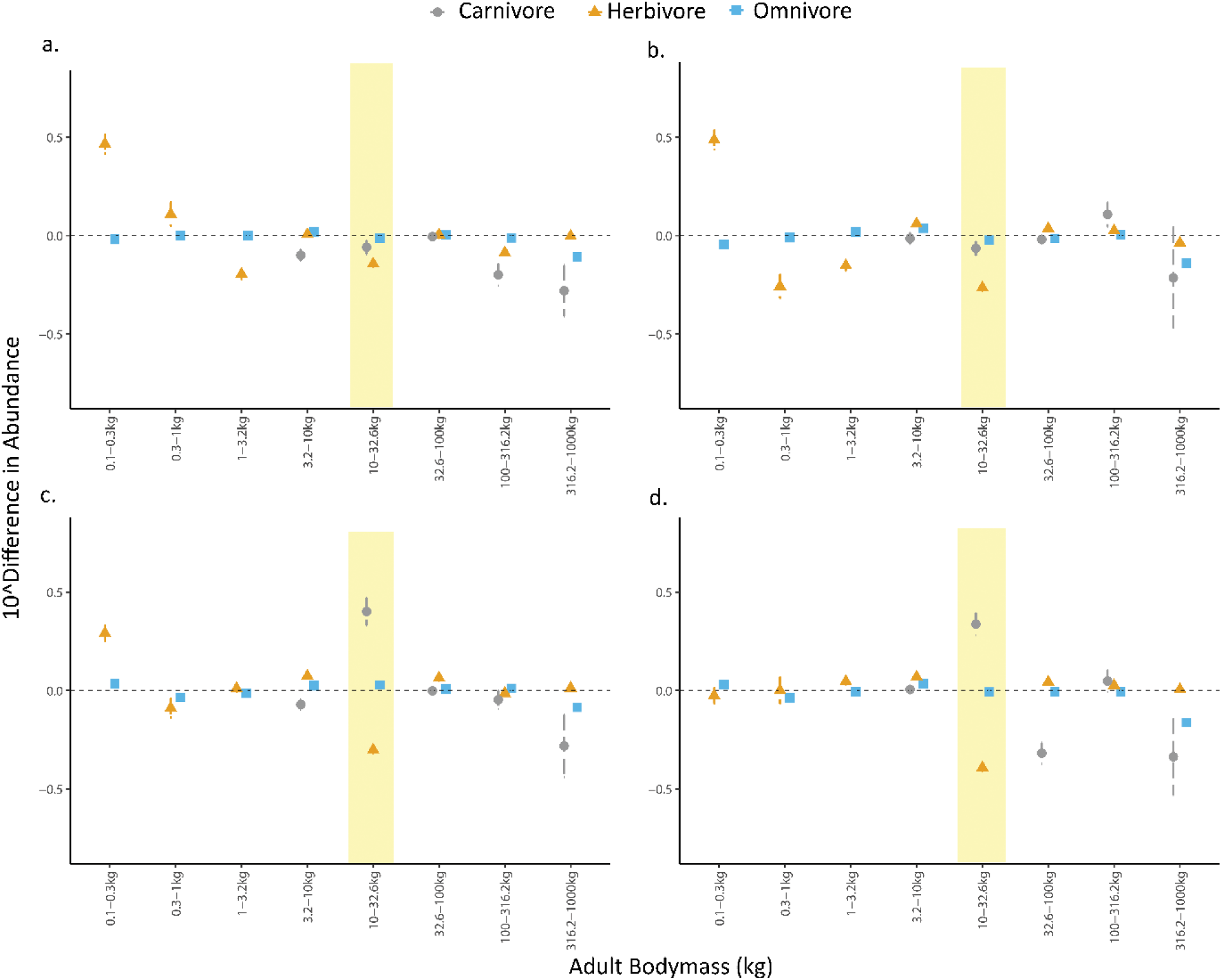
Changes in abundances (with 95% confidence intervals) of endothermic heterotrophs as a result of harvesting duiker-like herbivores (10-32.6kg group highlighted in yellow) at the rate of a. 20%, b. 50%, c. 70% and d. 90% of population year^-1^, by adult body mass. The horizontal dashed line indicates no significant impact of harvesting on abundances.

To account for temporal autocorrelation in animal abundances through time, changes in abundance due to harvesting were calculated as follows: change 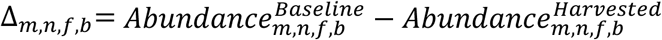, where abundances are measured in month *m*, for functional group *f* (herbivore/omnivore/carnivore) in body mass bin *b* in simulation *n*. For the purposes of this study, we compared total animal abundances without harvesting (‘Baseline’) to abundances where 20%, 50%, 70% and 90% of population was targeted (‘Harvested’). All data processing, statistical analysis and visualisation were done in R version 3.5.1 (R Core Team 2018), with some minor post-processing in Adobe Photoshop CS6.

## 3. Results

### 3.1 Validation

The probability of extinction, and the optimal harvesting rates expected yields from harvesting duiker-like herbivores predicted by the Madingley model were qualitatively and quantitatively similar to those predicted by the Beverton-Holt model, with a few notable differences.

Both models predicted a gradual decline in survival probability with increased harvesting (Figure 2a). Extinctions were noticeably more common without harvesting and at very low harvesting pressure in the Madingley model than in the Beverton-Holt model (at *φ*=0 survival probability of 0.86±0.13 and 0.99±0.001; 95% CI, respectively). The Beverton-Holt model also had a higher and a more pronounced threshold (at *φ* ≥0.15) harvesting rate above which extinction rate increased. The opposite was true at intermediate and high levels of harvesting, where survival rates were significantly higher in the Madingley than in the Beverton-Holt model. But both models estimated harvesting over 20% of population per year could result in a high risk of extinction.

In both models, expected yield was unimodal peaking at intermediate extraction rates (Figure 2b). Yields were maximised at an annual harvest rate of 20%-25% of the standing population. The interquartile ranges for yields did not overlap: the Madingley’s median yields were on average 11.67±1.49 (95% CI, *n*=30) times higher than the Beverton-Holt’s, and 4.64±0.44 (95% CI, *n* =30) times higher if mean yields were compared (Beverton-Holt’s yields were strongly right-skewed).

As a reminder, in the Madingley model, more than one species fell under our body-mass defined categorisation of duiker-like, and the model simulated an initial pristine ecosystem. Thus, a direct comparison between the Madingley and Beverton-Holt is not possible. For example, in addition to Peters’ and bay duiker, water chevrotain *Hyemoschus aquaticus* with mean body mass of 15kg, Ogilby’s duiker *Cephalophus ogilbyi*, 19.5kg, also fell into the duiker-like category. To at least help with the comparison, we added yields from harvesting bay duiker *C. dorsalis* to Peters’ duiker yields from the Beverton-Holt models, providing a lower bound on the predicted yield from multi-species harvesting. Now the difference between the two models fell by half: to 5.35±0.66 times for the median yields, and to 2.71±0.35 times for the mean yields. Given that the two modelling approaches are so very different (see Figure 1), we considered a match to within a factor of 5 to be sufficient to motivate further examination of the Madingley model predictions.

### 3.2 Ecosystem Impacts of harvesting duiker-like animals

At four levels of harvesting intensity (20%, 50%, 70% and 90%) of the duiker-like herbivores, there were a few general patterns in ecosystem responses to harvesting compared to the baseline (Figure 3). Harvesting above 20% of duiker-like population resulted in significant declines in duiker-like abundances (highlighted in yellow in Figure 3): on average, a 28% decline in duiker-like abundances was expected at *φ*=0.20, and a 59% decline in duiker-like abundances at *φ*=0.90.

The magnitude of the impact of harvesting on duiker-like abundances became clearer as we reduced the body mass bin ranges. When using the body mass range of 10-100kg, the duiker-like abundances declined by a factor of 2 (corresponding to differences in normalised abundances of 0.3) at *φ*=0.90 (the bold rectangle in Figure S1). When using the body mass range of 10-32.6kg, the duiker-like abundances declined by a factor of 2.5 (corresponding to differences in normalised abundances of 0.4) at *φ*=0.90 (the bold rectangle in Figure S2a). Finally, when using two even smaller body mass ranges of 10-17.8kg and 17.8-31.6kg, the duiker-like abundances declined by a factor of 3.2 (corresponding to differences in normalised abundances of 0.5) at *φ*=0.90 (the bold rectangle in Figure S2b). Interestingly, abundances of duiker-like herbivores with body masses of 17.8-31.6kg returned to pre-harvest levels in the last 10 years of harvesting (the bold rectangle in Figure S2b).

Harvesting duiker-like animals resulted in a number of changes in ecosystem structure. In particular, small-bodied (0.1-0.3kg) herbivores increased in abundance (by up to 206%) at low and medium-high levels of duiker harvesting (up to 70% of population year^-1^; Figure 3a-c), and remained unchanged at very high harvest rates (*φ*=0.90) (Figure 3d). Medium-sized (10-32.6kg) carnivores increased in abundance at high harvest rates (*φ*≥0.70). While large-bodied carnivores and omnivores (316-1000kg) were negatively affected by duiker-like harvesting, decreasing in abundance by between 37%-54% on average.

## 4. Discussion

Verifying and validating estimates of duiker harvesting from the multi-trophic Madingley model against a conventional Beverton-Holt model has shown high levels of quantitative and qualitative correspondence between these two independent models. Survival probability, estimated yields and maximum harvest rates were all comparable, despite differences in the focal harvested group, the model structure and other processes included. Both models estimated the optimal harvest rate at 20%-25% of the duiker population per year.

The Madingley model yields were around 5 times greater than the yield predicted by summing the Beverton-Holt models for the two duiker species (Figure 2). This difference is statistically different and obviously substantial in terms of the implied economic and nutrition value. However, we considered the result to be encouraging for several reasons. First, the sum over the Beverton-Holt models represents a lower bound on the predicted yields from single-species models, because it does not include other species that would fall into the same functional group and size range in reality, and in the Madingley model. Second, the Madingley simulations being with a truly pristine ecosystem, whereas the Beverton-Holt parameters are estimated from locations where humans have significantly impacted the ecosystems for thousands of years. Third, the Beverton-Holt predictions are themselves highly uncertain, reflecting parameter uncertainty (Barychka *et al.*, in prep.). Fourth, and most important, the Beverton-Holt predictions, and the Madingley model predictions, were generated from methodologies that are entirely independent, with the former employing a very simple model with species-specific parameters, and the latter employing a complex simulation model with climate as the only input. Given this major difference in methodology, we consider the good match in predictions for extinction and optimal harvest rate, and predicted yields within a factor of 5, to be highly encouraging.

The 10% extinction rate without harvesting in the Madingley (Figure 2a), which was not represented in the Beverton-Holt model, is arguably more realistic in reflecting the effects of environmental and demographic stochasticity that are absent in the Beverton-Holt (Lande *et al*., 1995; Lande *et al*., 1997; Bousquet *et al*., 2008). Although stochasticity could be easily added to a single-species model (Lande, 1998; Jonzén *et al.*, 2002), it emerges more realistically in the Madingley model as a result of interactions between and within trophic groups, and with their environment. Similarly, higher population persistence rates in the Madingley model than in the Beverton-Holt at moderate and high rates of harvesting were arguably more representative of real-life ecosystems, as: a) smaller animals would be more likely to avoid capture and reproduce (Wilkie and Finn, 1990), and b) predators would switch between similar-sized prey species as they became more rare (Allen, 1988). The population persistence dynamics revealed that keeping the risk of extinction below a maximum acceptable level of 10% on average (Mace and Lande, 1991) implied harvesting not more than 20% of duiker-like population year^-1^ - a rather low harvest rate, implying a trade-off that decision-makers may need to consider.

Here, the Madingley model was used to predict the effect of harvesting on ecosystem structure. Removing duiker-like herbivores had relatively low impacts on other functional groups, with the exception of small-bodied herbivores (which would likely compete with duikers for resources) and large-bodied predators. However, duiker-like herbivores contributed only between 2% and 4% of total abundance of similar-sized animals in the Madingley model, which could also explain this relatively low impact.

Studies of biological consequences of over-hunting on species in African tropical forest ecosystems generally focus only on the target species; declines in density were recorded in duikers and other mammals (e.g. Fitzgibbon, Mogaka and Fanshawe, 1995; Noss, 1998a; Gates, 1996). In terms of effects of removal of target species on non-target animal groups; in the Amazon, greater increases in abundances of large rodents and artiodactyls were reported in areas with higher levels of harvesting of arboreal monkeys, compared to moderately-hunted areas (Bodmer *et al.*, 1997). Very high abundances of common opossums *Didelphis marsupialis* and spiny rats *Proechimys* spp. were reported in heavily fragmented forests of Brazil and central Panama, explained by the absence of their predators and/or competitors (Adler, 1996; da Fonseca and Robinson, 1990). Fa and Brown (2009) predicted that the abundance of non-target small and medium-sized species could remain unchanged or even increase depending on the availability of their prey and removal of competitors and other predators. According to Wright (2003), large-bodied species preferred by hunters would decline with harvesting pressure; the less desirable species would first increase due to lower competition for resources, and then decline; and small untargeted species would increase steadily. The trophic cascades theory predicts that higher abundances of mid-level consumers should result in lower abundance of basal producers (assuming ‘top-down’ control) (Pace *et al.*, 1999; Kennedy, 2012; Palmer *et al.*, 2015). However, changes in higher trophic levels do not always propagate to lower levels or have significant ecosystem impacts; higher resilience to perturbations is possible in systems with high trophic diversity and complex food webs (Wright, 2003; Pace *et al.*, 1999).

From the point of view of a bushmeat manager considering the wider ecosystem impacts of harvesting, the system, as indicated by the Madingley model, was relatively robust to intensive harvesting. Many animals were heavily depleted but did not become extinct, smaller-bodied animals increased in abundance, and vacant ecological niches were being quickly filled-in by, presumably, more resilient faster-reproducing animals (Adler, 1996; da Fonseca and Robinson, 1990). However, harvesting intensively also resulted in a very different ecosystem structure (Scheffer *et al.*, 2001; Barychka, Mace and Purves, 2019), dominated by small-bodied short-lived animals. Considering the trade-off between high yields now, and lower yields, lower species diversity, and a different ecosystem structure and functioning later, should be a part of decision-making process in bushmeat management.

Our harvesting protocol was relatively simple. Harvesting was applied to a single location approximately 100km x 100km; no inter-cell migration was allowed. Although duiker home ranges are relatively small, around 0.10km^-2^ (Payne, 1992), in reality local duiker populations would likely disperse (depending on strength of pressure on neighbouring ecosystems) and therefore replenish nearby areas, most likely then increasing species overall tolerance to pressure (Fa and Brown, 2009). We assumed constant non-adaptive harvesting which was not affected by the return per unit effort, the selectivity of hunters (Wright, 2003), or any other socioeconomic factors such as proximity to roads or access to salaried employment (Nielsen, 2006; Nielsen, Jacobsen and Thorsen, 2014). No provision was made in the model for the potential wastage due to animals captured and discarded as unsuitable for sale or consumption, or animals escaping after being injured (and likely dying later on), though it could add a quarter to recorded harvesting mortality (Noss, 1998a).

The Madingley model’s main strengths are its generality and ability to look at any functional species group and location, including ones that have not yet been studied in any detail and thus are lacking in data (Purves *et al.*, 2013; Bartlett *et al.*, 2016). Here, we used the model against one of the most common and best studied bushmeat species, so that we could compare the Madingley model results to those from traditional methods. The Madingley model was able to produce reasonable estimates for duiker-like harvesting dynamics based solely on climate data and given ecological processes. While the Beverton-Holt model was able to capture the salient features of single-species harvesting (Lande, Sæther and Engen, 1997; Fryxell *et al.*, 2010), in the absence of population parameter estimates the Madingley model could offer adequate indication of harvesting outcomes. Therefore, the main value may come from using the Madingley model (or models like it) for location and species that have been barely studied at all.

Moreover, there is a lack of understanding of synergies and interactions within ecosystems (da Fonseca and Robinson, 1990; Wright, 2003) which we may not be able to address using traditional modelling for some time. Predicting dynamics and potential impacts of multi-species harvesting has not been considered feasible for many real-life populations (Hooper *et al.*, 2005). These results suggest that in the absence of well-informed empirical models mechanistic models such as the Madingley General Ecosystem Model could provide helpful approximations of such dynamics.

## Author’s contributions

TB, GMM and DP conceived of the ideas and designed methodology; TB ran computer simulations, analysed data and lead the writing of the manuscript. All authors contributed critically to the drafts and gave final approval for publication.

## Acknowledgements

This work was funded by the Natural Environment Research Council, UK.

