## Supplement Figures S1 and S2 for "The Madingley General Ecosystem Model predicts bushmeat yields, species extinction rates and ecosystem-level impacts of bushmeat harvesting"

Figure S1. Total abundances of carnivores, herbivores and omnivores (on log-scale, normalised to month 0; with 95% confidence intervals) over 30 years, without (grey) and with (orange) harvesting of 90% of duiker-like herbivores. Animals were grouped into body mass bins. The impact of harvesting is explored under increasingly high resolution, by reducing the sizes of the body mass bins from Figure S1 to Figure S2a, to Figure S2b. The target group (duiker-like herbivores) is emphasized by the bold rectangle; arrows and dotted rectangle indicate animal groups which were inspected in more detail in Figure S2a and 2b.

Figure S2. Total abundances of carnivores, herbivores and omnivores (on log-scale, normalised to month 0; with 95% confidence intervals) over 30 years, without (grey) and with (orange) harvesting of 90% of duiker-like herbivores. The impact of harvesting is explored under increasingly high resolution, by reducing the sizes of the body mass bins from a. to b. The targeted group (duiker-like herbivores) is emphasized by the bold rectangle. Arrows and the dotted rectangle in a. indicate animal groups that were inspected in more detail in b.

Figure S1

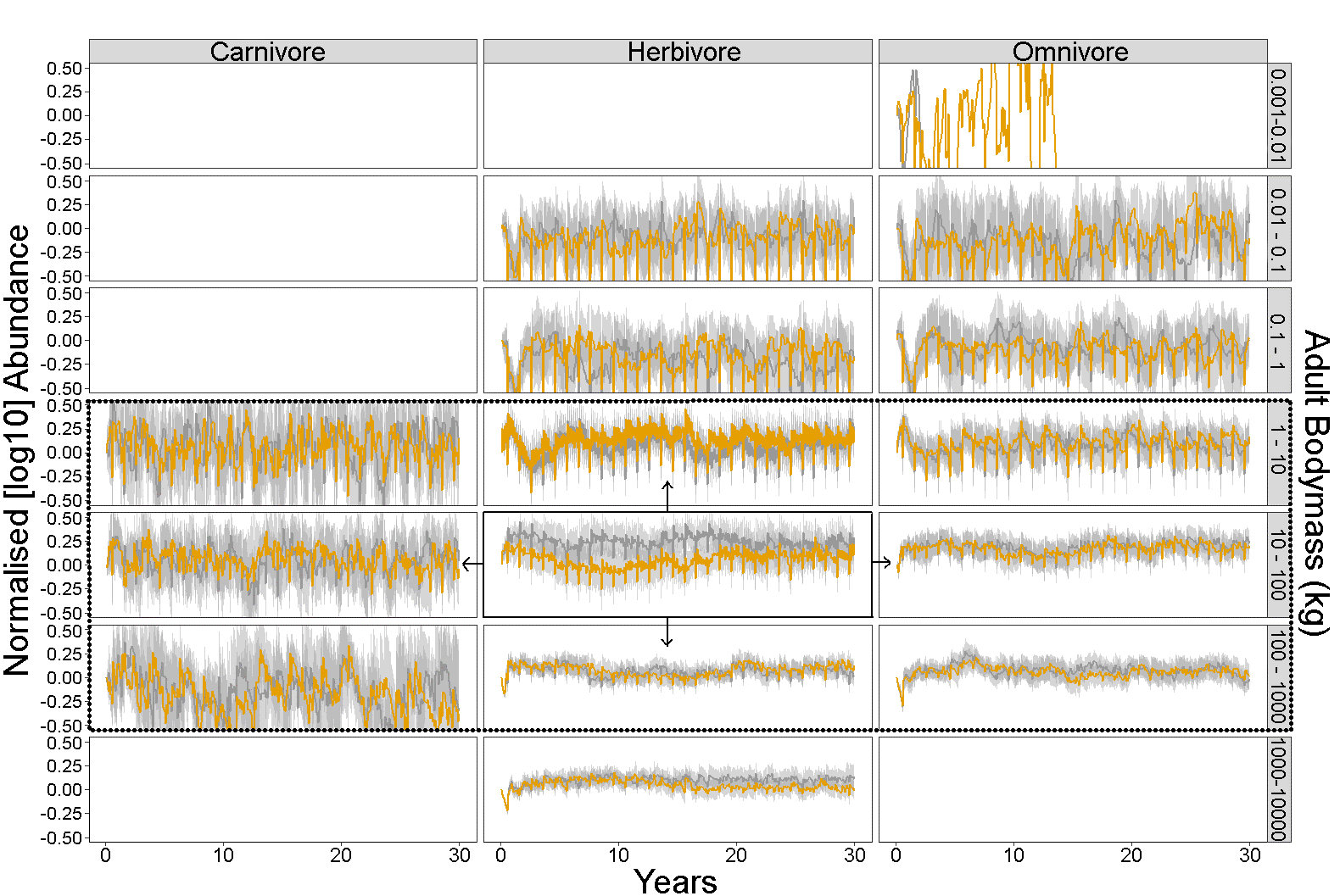

Figure S2

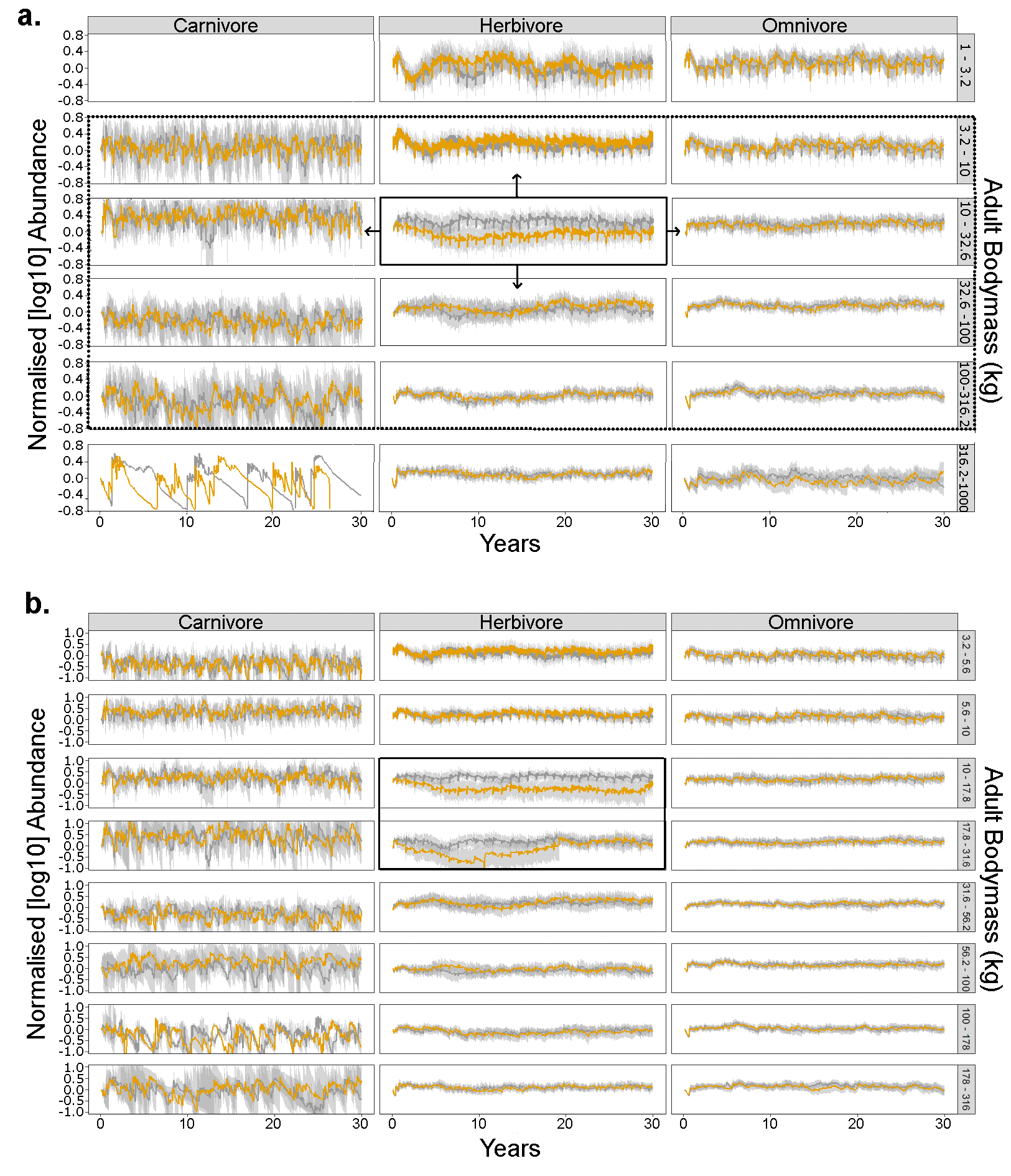
